# Combining Information from Crosslinks and Monolinks in the Modelling of Protein Structures

**DOI:** 10.1101/2020.03.25.007104

**Authors:** M. Sinnott, S. Malhotra, M.S. Madhusudhan, K. Thalassinos, M. Topf

## Abstract

Monolinks are produced in a Chemical Crosslinking Mass Spectrometry experiment and are more abundant than crosslinks. They convey residue exposure information, but so far have not been used in the modelling of protein structures. Here we present the Monolink Depth Score (MoDS), for assessing structural models based on the depth of monolinked residues, corresponding to their distance to the nearest bulk water. Using simulated and reprocessed experimental data from the Proteomic Identification Database, we compare the performance of MoDS to MNXL - our previously-developed score for assessing models based on crosslinking data. Our results show that MoDS can be used to effectively score model structures based on monolinks, and that combining it with MNXL leads to overall higher scoring performance. The work strongly supports the use of monolink data in the context of integrative structure determination. We also present XLM-Tools, a programme to assist in this effort, available at: https://github.com/Topf-Lab/XLM-Tools.

## Introduction

The determination of protein structure is essential for our understanding of the processes governing biological functions. The most common techniques for determining protein structures are X-ray crystallography, NMR spectroscopy and cryo-electron microscopy. However, these techniques can be time-consuming and are not always useful due to factors such as protein size and flexibility. An alternative approach is computational prediction of protein structure, using various approaches, such as homology modelling, threading and *ab-initio* (Deng et al., 2018). The quality of the predicted models varies depending on both the modelling method employed and the system being investigated. A challenge (in the field) is to rank models according to their accuracy (Won et al., 2019).

Chemical Crosslinking Mass Spectrometry (XLMS) is a technique which can assist in predicting monomeric structural models (Matthew Allen Bullock et al., 2016; Young et al., 2000; Zelter et al., 2010) as well as for multi-subunit protein complexes (Bullock et al., 2018a; Kosinski et al., 2015; Tosi et al., 2013; Wu et al., 2012) and has been gaining popularity in recent years. It is a technique that allows the generation of solvent accessible distance restraints for structural modelling purposes (Jan Seebacher et al., 2006; Leitner et al., 2010). It has advantages over the traditional structural determination methods such as being relatively quick and cheap to perform, can be performed in samples that are not entirely pure, and recently has been used in the context of whole cellular environments (Kaake et al., 2014).

XLMS is performed by subjecting a protein or a complex to a chemical crosslinking reagent by covalent labelling, which will then react with solvent accessible residues on the protein surface (the type of residue depends on the chemistry of the crosslinking reagent). The two residues may be crosslinked if they are within a maximum Solvent Accessible Surface Distance (SASD) of each other (dictated by the length of the crosslinker). SASD is the shortest distance between the two residues on the surface of the protein (Matthew Allen Bullock et al., 2016). After crosslinking, the protein is digested using a protease (normally trypsin), yielding four different types of peptides: unmodified, looplinks, crosslinks and monolinks. At this point, a peptide enrichment procedure (usually strong cation exchange) is often employed to remove unmodified peptides, which are of no informational value (Leitner et al., 2010). Looplinked, crosslinked and monolinked peptides are then sequenced using LC-MS/MS to identify the residues which have been modified.

Crosslinks are the most information rich peptides in an XLMS experiment. They provide two types of complementary information: they are a marker of solvent accessibility for the two residues associated and they also provide a distance constraint between the two. We recently utilised this information to produce the Matched and Non-Accessible Crosslink Score (MNXL) (Matthew Allen Bullock et al., 2016). This score, which encapsulates both the distance and solvent accessibility information in a crosslink, showed increased performance compared to previously-used scores that are based only on the Eucledian distance (Herzog et al., 2012; Leitner et al., 2012). The MNXL score compares a list of SASDs between the crosslinkable residues against a list of experimentally determined crosslinks. The SASDs provide information on each potential crosslink in terms of solvent accessibility and crosslink length (Matthew Allen Bullock et al., 2016). The MNXL program (i) will check whether an input BS3/DSS crosslink is solvent accessible (number of non-accessible crosslinks, NoNA); (ii) will check if the length of the crosslink is within a maximum bound of 33 Å (NoV), and (iii) will score the solvent accessible crosslinks according to a pre-determined probability distribution of experimentally determined crosslinks (Expected Solvent Accessible Surface Distance – ExSASD). In the case of MNXL, SASDs are generated using a program called Jwalk (Matthew Allen Bullock et al., 2016). MNXL was benchmarked on data generated with BS3/DSS crosslinks, both of which have a linker length of 11.4 Å.

Investigating the utility of solvent accessibility for scoring models based on crosslinks provides an opportunity to explore another type of information coming from an XLMS experiment, namely monolink information. A monolink can be thought of very simply as a covalent label showing residue exposure. The advantage of monolinks comes from their higher frequency identification than crosslinks in an XLMS experiment. The recovery of crosslinks has been estimated to be between 8-25% (Herzog et al., 2012) in previous experiments. For monolinks, however, although recovery estimates have not been published, it is presumed to be higher, and in the range of 50-70%.

Here, we introduce the Monolink Depth Score (MoDS) as a scoring function to assess a model structure based on this chemical labelling information. With MoDS we utilise residue depth, a measure which calculates the distance of a residue from the nearest bulk water molecules (Tan et al., 2011), therefore excluding non-bulk water found in cavities and trapped inside the protein structure. We first show the utility of this score on a large benchmark with simulated monolinks. Next, we demonstrate the beneficial effect of combining monolink and crosslink-based scores, which we name Crosslink-Monolink (XLMO) score. We also assess the score on a smaller benchmark based on publicly available experimental XLMS data.

## RESULTS

### Theory

The Monolink Depth Score (MoDS) is described as:

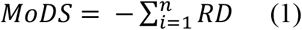

Where *i* is every amino acid in the protein which has an associated monolink and residue depth is the Residue Depth of that amino acid. *n* is the total number of monolinked residues in the protein.

The input to the method is a list of amino acids which have been modified by a crosslinker and a list of the residue depths calculated by DEPTH from the protein structure (Tan et al., 2011). It will then remove all non-bulk waters, where the latter are defined as 4 or more water molecules in a 4.2 Å radius (Figure 1A). Non-bulk waters are usually located in cavities and trapped inside the protein structure, and therefore are not representative of the solvent environment of the protein. The solvent configurations are then sampled in 25 different orientations of the protein (solvation cycles), and residue depth is then measured as the average distance of the residue to the nearest bulk water molecule over all solvation cycles. Other than defining the minimum number of “surviving” solvent as 4, default settings were used in DEPTH calculations (Tan et al., 2011).

**Figure 1:**
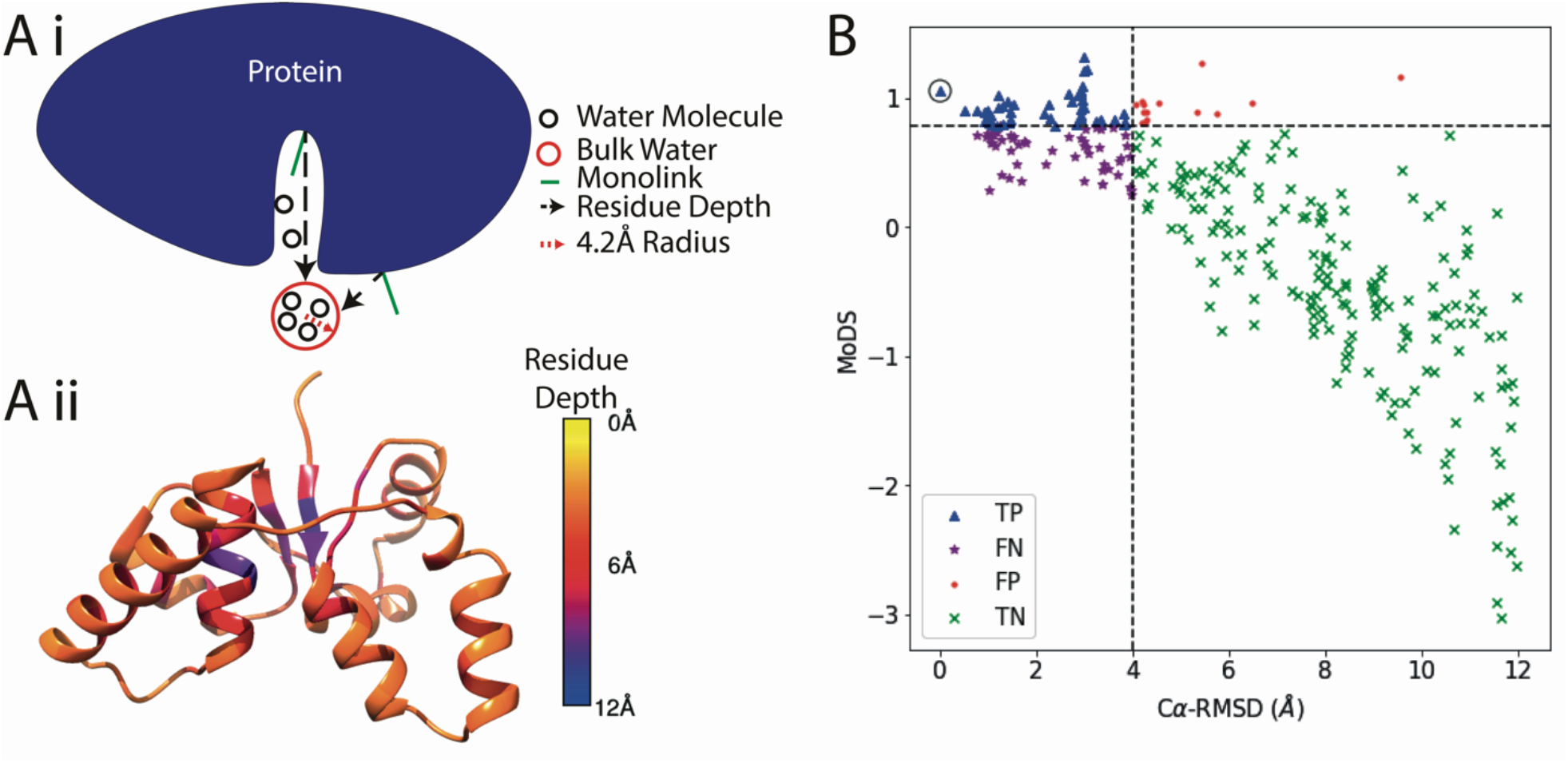
Illustration of Monolink Depth Score (MoDS). A) i. Schematic of the measurement of residue depth, which is a precursor step to MoDS calculation. The shortest possible distance is calculated between the nearest bulk water and the residue in question, ignoring any non-bulk water molecules. ii. Structure of 3H87:A coloured by residue depth. B) Scatter plot of MoDS *vs.* Cα-RMSD for a target from the simulated benchmark (3H87:A). Points are split into True Positive (TP), False Positive (FP), False Negative (FN) and True Negative (TN). The target structure (which is ranked within the top 10 scoring models) is circled.

We also combine MoDS with information from crosslinks using the Matched and Non-Accessible Crosslink (MNXL) score (as described in the introduction) (Matthew Allen Bullock et al., 2016). We calculate a Crosslink Monolink score (XLMO) by combining these two scores. We do this by calculating the z-scores for MoDS and MNXL and then adding these.

### MoDS, MNXL and XLMO performance on the simulated benchmark

To evaluate MoDS, MNXL and XLMO performance statistically we tested it on the 3DRobot benchmark downloaded from the Zhang Lab (Deng et al., 2016), a benchmark of 200 model structures each of which has 300 decoy model structures. We simulated crosslinks and monolinks on these models at 20% recovery for crosslinks and 50% for monolinks (see STAR Methods).

Figure 1B shows the results from chain A of a target with PDB Id: 3H87 from the simulated benchmark (3H87:A), with monolinks simulated at 50% recovery. In this example, the simulated monolinks are a good predictor of structure, owing to a correlation of 0.85 between MoDS and Cα-RMSD and precision and AUC of 0.92 and 0.94, respectively (Table S2).

We compared the performance of the three scores MoDS, MNXL and XLMO at a constant recovery rate of 20% for crosslinks and 50% for monolinks over the entire benchmark (Figure 2). Density plots representing the different scores in terms of AUC, correlation and precision (Figure 2A) show that MNXL performs better than MoDS over all statistical measures (significant with t-test P ≤ 0.05). XLMO on average outperforms both MNXL and MoDS in terms of AUC (0.82, 0.78, 0.77, respectively); Spearman Correlation (0.59, 0.52, 0.48, respectively) and Precision (0.67, 0.64, 0.54, respectively) (Table S2) (significant with t-test P ≤ 0.05 for all measures).

**Figure 2:**
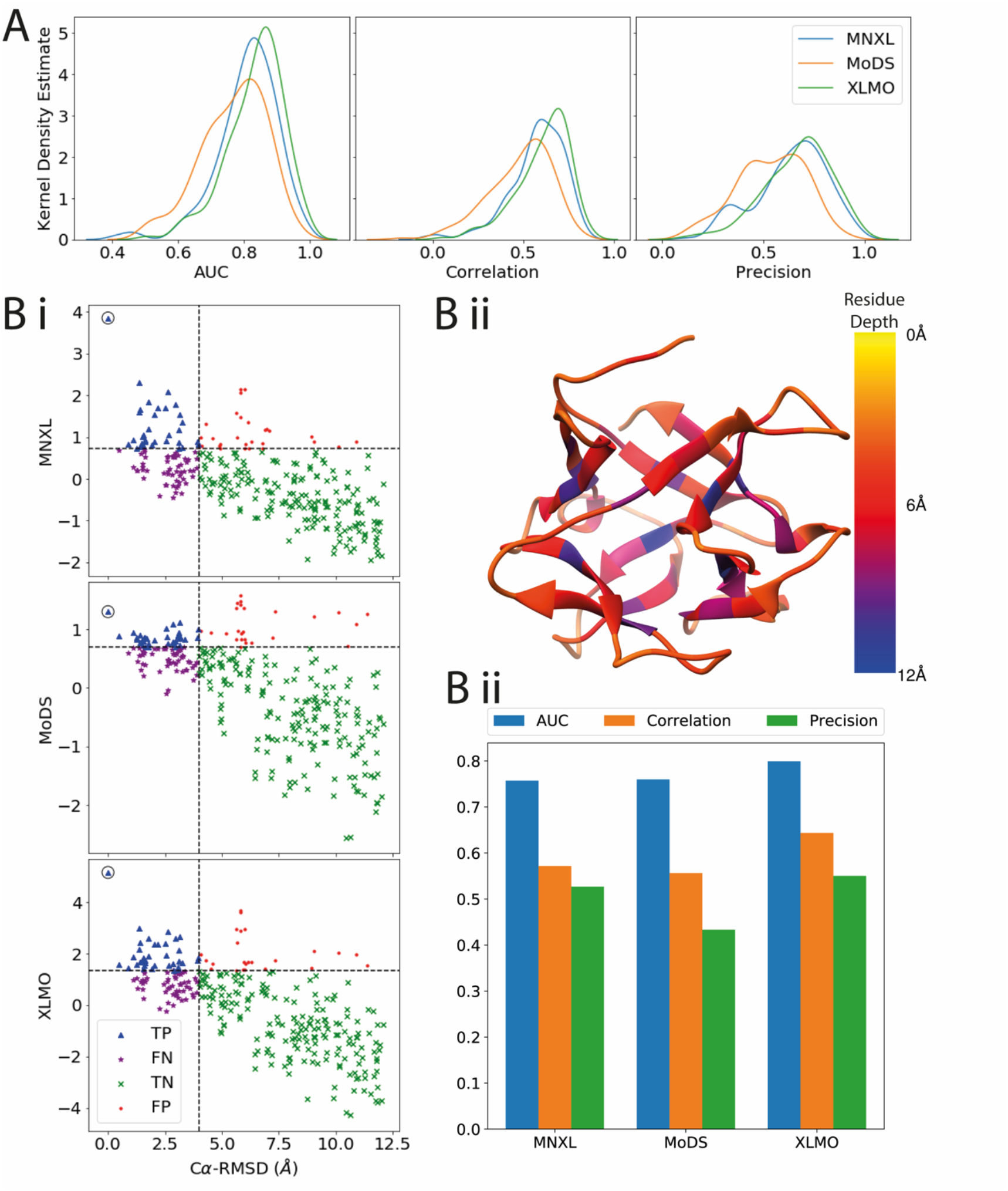
MoDS, MNXL and XLMO scoring for the simulated benchmark. A) Kernel Density Estimate Plots comparing the different scoring methods in terms of correlation, precision and AUC. Crosslinks are simulated at 20% recovery, monolinks at 60%. B) i. Scatter plots of each scoring methods *vs.* Cα-RMSD for 4IZX:A. Points are split into True Positive (TP), False Positive (FP), False Negative (FN) and True Negative (TN) and the target structure is circled. ii) Structure of 4IZX:A coloured by residue depth. iii) Bar plot shows statistics for MNXL, MoDS and XLMO.

4IZX:A, which is a *Macrolepiota proceraricin* B-like lectin ß-trefoil structure is an example which demonstrates the improved performance of XLMO (Figure 2bi). Its residue depth varies greatly across the residues in the target structure (Figure 2Bii). MoDS shows reduced performance compared to MNXL in this model (Figure 2biii), however, combining these two scores (in XLMO) exhibits an increase in performance by all statistical measures (Table S2).

### MoDS, MNXL and XLMO performance on the experimental dataset

A smaller dataset containing experimental crosslink data was used to assess the performance of the scores with experimentally-determined crosslinking results (Table 1). These datasets were acquired from the Proteomic Identification Database (PRIDE) and reprocessed (see STAR methods). The average residue depth for both crosslinked and monolinked residues over the entire dataset is around 6.35 Å (± 2.55 Å standard deviation). The values show overall worse performance than the simulated benchmark, with average AUC of 0.71 for MNXL, 0.71 XLMO and 0.73 for MoDS (Table 2). Similar to the simulated benchmark, the mean scores here also show that MoDS performs better than MNXL in most cases, with the exceptions being CASP targets Human Exonuclease V, X0975, and Target Surface Protein from *Enterococcus faecalis,* X0987. XLMO performance is varied in comparison to MoDS and MNXL over the benchmark, it performs better in 0/6 models by AUC (performs the same as MoDS for 3/6), 2/6 models in terms of correlation and in 1/6 in terms of precision (Table 2, excluding X0987:D1 and D2, which will be discussed later). Due to the small size of the benchmark, we could not show a significant difference between the scores in this benchmark.

**Table 1:**
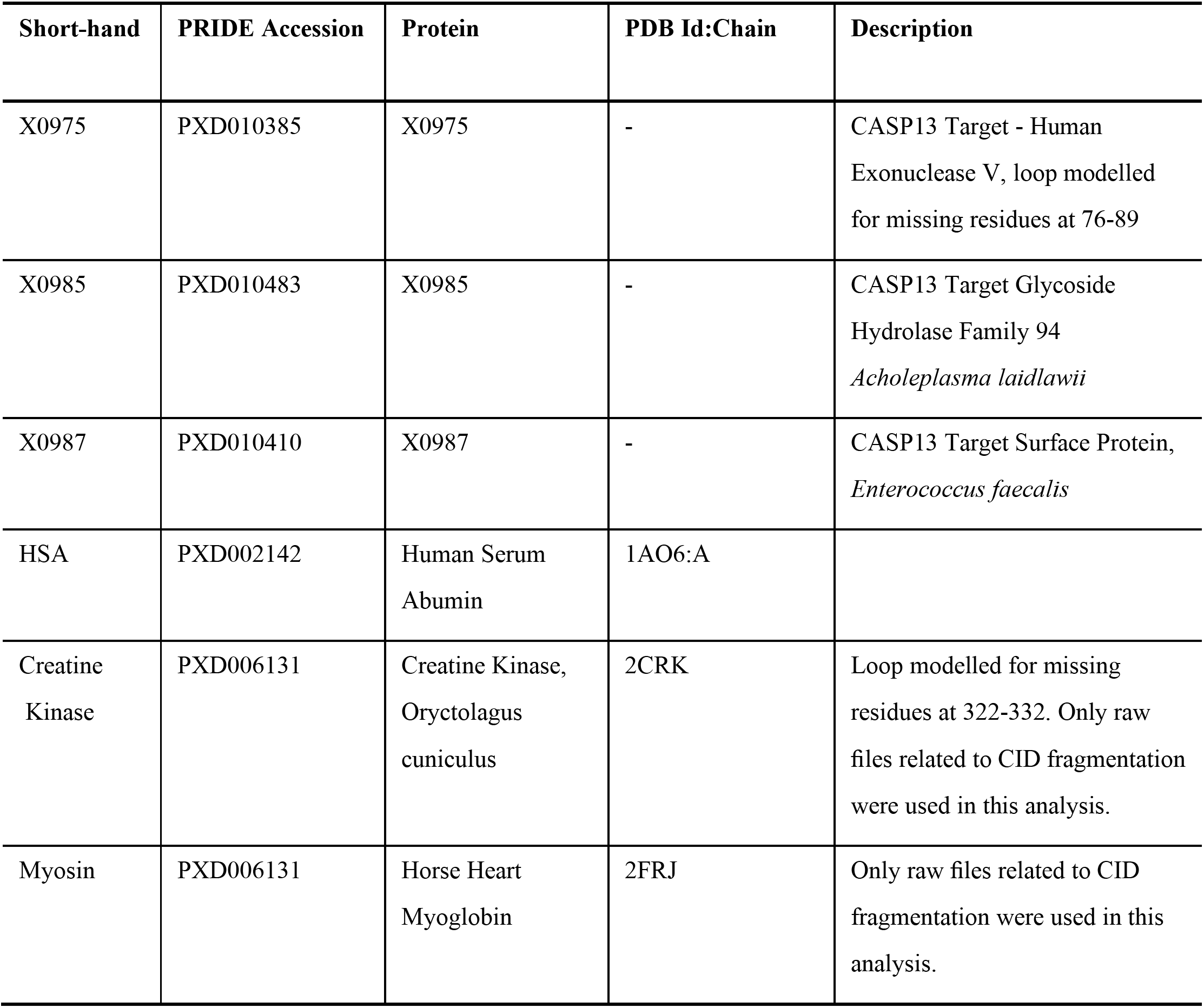
Experimental XLMS benchmark used in this analysis and PDB structures which they are associated to.

| Short-hand | PRIDE Accession | Protein | PDB Id:Chain | Description |
| --- | --- | --- | --- | --- |
| X0975 | PXD010385 | X0975 | - | CASP13 Target - Human Exonuclease V, loop modelled for missing residues at 76-89 |
| X0985 | PXD010483 | X0985 | - | CASP13 Target Glycoside Hydrolase Family 94<br>*Acholeplasma laidlawii* |
| X0987 | PXD010410 | X0987 | - | CASP13 Target Surface Protein,<br>*Enterococcus faecalis* |
| HSA | PXD002142 | Human Serum Albumin | 1AO6:A |  |
| Creatine Kinase | PXD006131 | Creatine Kinase, *Oryctolagus cuniculus* | 2CRK | Loop modelled for missing residues at 322-332. Only raw files related to CID fragmentation were used in this analysis. |
| Myosin | PXD006131 | Horse Heart Myoglobin | 2FRJ | Only raw files related to CID fragmentation were used in this analysis. |

**Table 2:** Statistics from the experimental benchmark. Mean and Standard Deviation (STD) values exclude the two domains of X0987 (D1 and D2).

| Model | AUC |  |  | Correlation |  |  | Precision |  |  |
| --- | --- | --- | --- | --- | --- | --- | --- | --- | --- |
|  | MNXL | MoDS | XLMO | MNXL | MoDS | XLMO | MNXL | MoDS | XLMO |
| *HSA* | 0.71 | 0.82 | 0.81 | 0.29 | 0.60 | 0.53 | 0.38 | 0.48 | 0.52 |
| *Creatine Kinase* | 0.81 | 0.86 | 0.86 | 0.54 | 0.64 | 0.65 | 0.42 | 0.47 | 0.45 |
| *Myosin* | 0.61 | 0.72 | 0.68 | 0.17 | 0.38 | 0.30 | 0.38 | 0.60 | 0.52 |
| *X0975* | 0.50 | 0.50 | 0.50 | 0.04 | 0.00 | 0.01 | 0.10 | 0.07 | 0.08 |
| *X0985* | 0.80 | 0.84 | 0.84 | 0.49 | 0.45 | 0.51 | 0.33 | 0.40 | 0.35 |
| *X0987* | 0.81 | 0.51 | 0.70 | 0.43 | -0.14 | 0.18 | 0.42 | 0.10 | 0.32 |
| *X0987:D1* | 0.60 | 0.71 | 0.68 | 0.04 | 0.43 | 0.28 | 0.37 | 0.42 | 0.42 |
| *X0987:D2* | 0.76 | 0.72 | 0.77 | 0.35 | 0.22 | 0.33 | 0.30 | 0.32 | 0.33 |
| *Mean*<br>*(Without*<br>*X0987 D1*<br>*and D2)* | 0.71 | 0.71 | 0.73 | 0.33 | 0.32 | 0.36 | 0.34 | 0.35 | 0.37 |
| *STD*<br>*(Without*<br>*X0987 D1*<br>*and D2)* | 0.12 | 0.15 | 0.12 | 0.18 | 0.29 | 0.22 | 0.11 | 0.20 | 0.15 |

Creatine kinase (Figure 3) has only 9 crosslinks but 36 monolinks, and this is reflected in the higher scoring for MoDS over MNXL (residue depth for crosslinks is also more variable for crosslinks than monolinks, Figure S3). However, XLMO has a similar performance to MoDS for AUC and correlation (0.86 and 0.65), but a slight fall in precision (0.47 vs 0.45). On the other hand, in the example of CASP target, Glycoside Hydrolase Family 94 from *Acholeplasma laidlawii* X0985 (Figure 3), the two-domain protein has a larger number of crosslinks (59 crosslinks) relative to monolinks (68 monolinks). Surprisingly, for this model (bearing in mind the high number of crosslinks), MoDS outperforms MNXL for all statistical measures. However, combining the information from both crosslinks and monolinks (in XLMO) results in a performance decrease for precision, no change in AUC and a small increase in correlation (0.51 for XLMO vs 0.45 for MoDS, Table 2). Another example of a two-domain protein is the CASP Target Surface Protein, *Enterococcus faecalis,* X0987, which performs much worse with MoDS scoring (Figure 3). It has 44 monolinks vs. 49 crosslinks. This large number of crosslinks results in good MNXL performance (AUC: 0.81, Correlation 0.43 and Precision 0.42, Table 2). However, MoDS performance is much lower, with an AUC of 0.51, a correlation of −0.14 and a precision of 0.1. This poor MoDS performance affects XLMO performance, which is not improving over MNXL in this case (see more details below).

**Figure 3:**
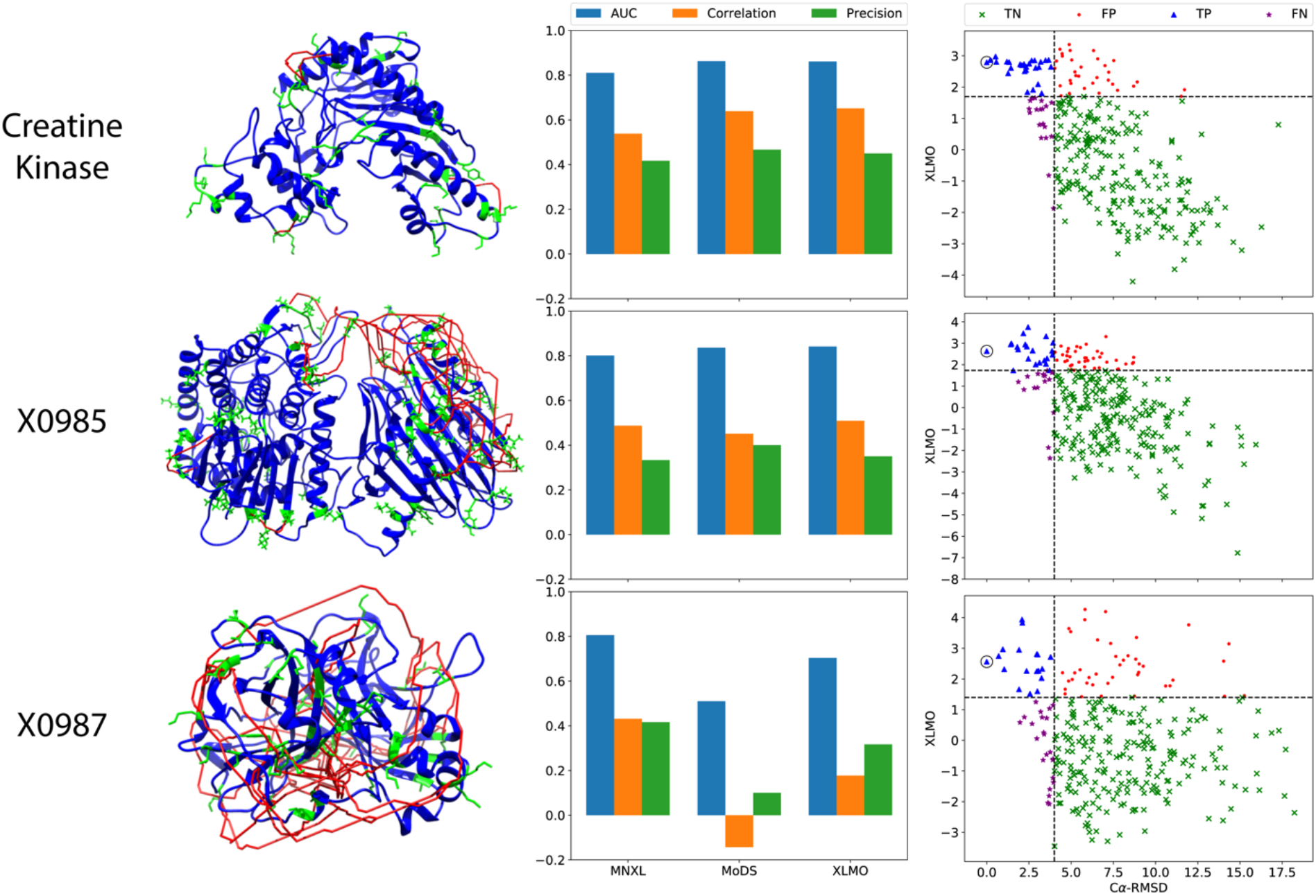
Examples from the experimental dataset (CK for Creatine Kinase, and CASP13 targets X0985 and X0987 are CASP13 targets). Crosslinks shown on corresponding structures (left) coloured red, monolinked residues shown coloured in green. Bar graphs (middle) show statistics for MNXL, MoDS and XLMO. Scatter plots (right) show the 300 models for each protein scored with XLMO against Cα-RMSD.

### Effect of residue depth on performance

To investigate the reasons for large differences in the experimental benchmark performance, we observed the differences in residue depth for all monolinked residues between the target structure and the top 5-ranked models in terms of MoDS for Creatine Kinase (well performing) and X0987 (poorly performing) (Figure S3). For Creatine Kinase only 10 residues differ by more than 0.5 Å or less than −0.5 Å in residue depth from the target structure. Most of these residues are located in a flexible and exposed region of the protein towards the C-terminal end.

This is in contrast with X0987, where surprisingly many of the residues which have associated monolinks have high residue depth in the target structure, ranging between 4.06-15.29Å. In the top-ranked models these residues are up to 8 Å shallower than in the target structure in terms residue depth. Looking in detail into this case, we observed that the performance of MoDS on domain 1 (D1, residues 11-195) and domain 2 (D2, residues 196-402) alone exhibits a sizeable increase as compared to the full protein (precision for MoDS increases from 0.10 for the full X0987 target structure to 0.42 for X0987:D1 and 0.32 for X0987:D2, Table 2). The reason is likely that in the full structure some of the monolinks in D2 fall into the interface region between the two domains, therefore exhibiting high residue depth (Figures S3). This results in high variance for residue depth of monolinks in the overall structure and poor overall performance. However, when separated, these residues in D2 have a lower residue depth (Figure S3) and scoring is improved for this domain.

### Effect of recovery on model scoring performance

The recovery analysis of the entire simulated benchmark (Figure 4) shows that the MoDS performance does not plataeu, and continues to rise to 100% recovery. At the 50% recovery point (marking the assumed monolink recovery) MoDS has a correlation of 0.48 with Cα RMSD, Precision of 0.54, and AUC of 0.77. XLMO performance also rises to 100% but at a lower rate than MoDS, although MoDS performance never overtakes XLMO. At the 50% recovery point XLMO has a correlation of 0.60 with Cα RMSD, Precision of 0.66, and AUC of 0.83.

**Figure 4:**
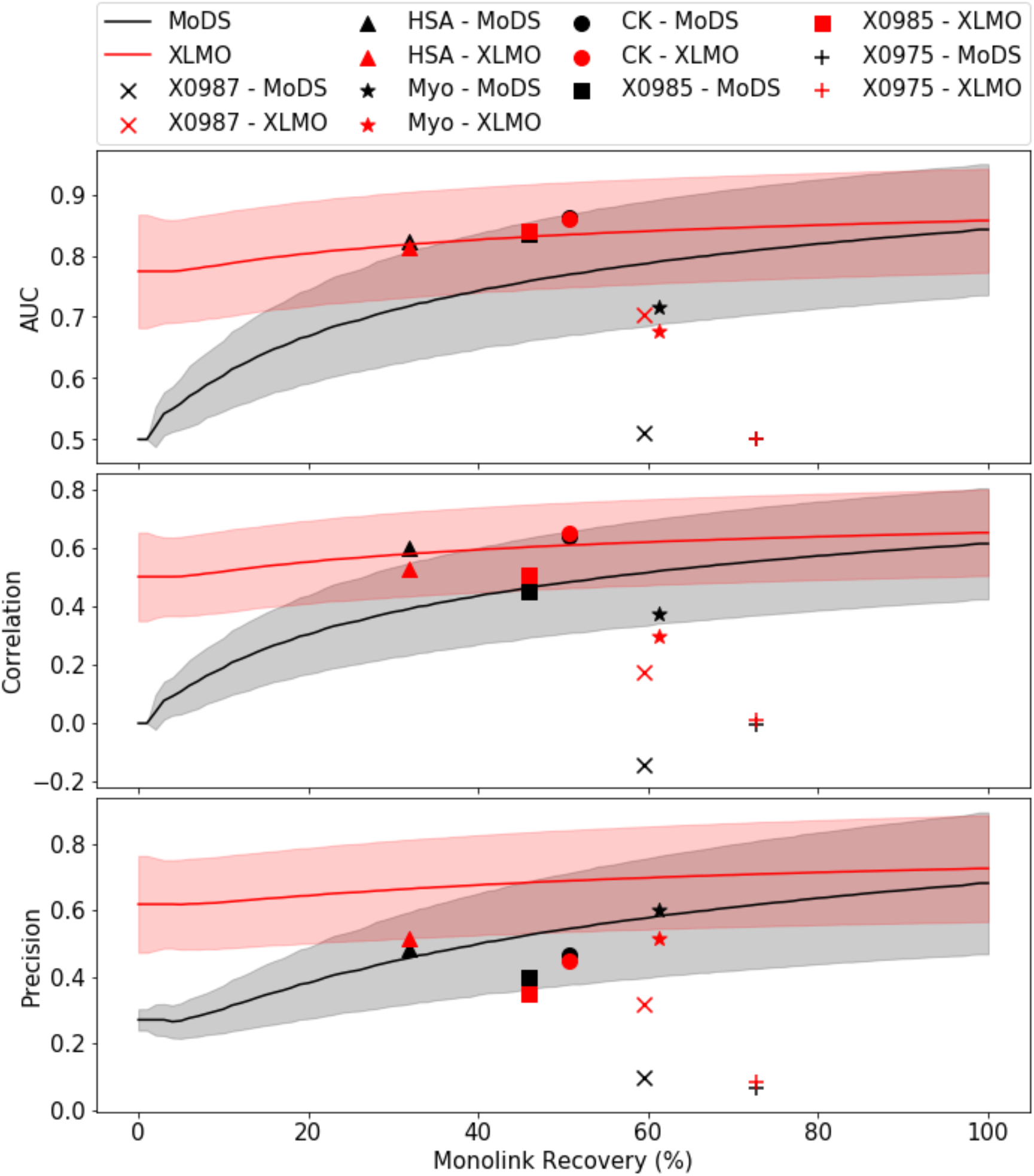
Recovery analysis of the simulated dataset with points from the experimental dataset (based on MoDS and XLMO scoring in black and red, respectively). Performance for MoDS and XLMO is shown overlaid. Solid line denotes mean score over the entire benchmark, shaded area shows standard deviation around the mean. Performance is shown as AUC, Spearman Correlation Coefficient (Correlation) and Precision, denoting the relationship between Ca-RMSD and MoDS/XLMO.

As shown previously, the performance of experimental models varies greatly (Figure 4). Cases Creatine Kinase, human serum albumin (HSA), Myosin and X0985 perform in the range of what would be expected within the error range of the simulated benchmark. However, other cases appear to underperform given their monolink recovery rate, showing no obvious correlation between the recovery rate and the performance for either MoDS or XLMO. For example, X0975 has a 60% monolink recovery rate but appears to exhibit poorer performance than expected.

## DISCUSSION

The information presented in this paper shows that monolinks can be used to obtain structural information on protein structures. Our analysis focused mainly on single domains but we also show that it can work on chains containing two domains (e.g. X0985). The monolink information is additional to the crosslink data coming from XLMS experiments that is typically used in the modelling of protein structure. Using simulated and experimental benchmarks, we showed that our monolink-based score (MoDS) carries valuable information and in a few cases performs better than the crosslink-based score (MNXL (Matthew Allen Bullock et al., 2016)). Furthermore, combining the two scores in the XLMO score can lead to improved assessment of protein structure models when monolinks are simulated. For the case of experimental monolinks, however, there is more variability in the score’s performance. We have looked into the possibility of applying different relative weights to the MNXL and MoDS in the XLMO score; however, no significant difference in the results of the simulated benchmark have been observed.

The recovery analysis shows that MoDS continues to rise in performance to 100% while the combined score (XLMO) increases at a slower rate, likely due to the influence of the MNXL score (which typically plateaus at around 25% crosslink recovery rate (Matthew Allen Bullock et al., 2016). However, there is a great difference in performance between MoDS and XLMO, with the latter performing a lot better especially at low monolink recovery rates.

Although mean scores for the experimental benchmark indicate a similar picture in the experimental dataset, none of the models show an overall improvement for XLMO over all statistical measures. XMLO appears to perform better in cases where poor scoring is achieved for either MNXL or MoDS. XLMO can therefore balance out this poor performance obtained from individual scores.

### Relevance of Simulated Monolinks

The simulation of monolinks was performed in order to obtain a realistic scoring performance, by randomly resampling crosslinks at 20% recovery and monolinks at 50% recovery. This approach allows all possible monolinks to be formed, thereby have maximal spatial coverage over the protein surface. As a result, simulated monolinks are likely to perform better on average than experimental monolinks for a given recovery rate.

Prior to this study we could not find a reported recovery rate estimation for monolinks in the literature. Our small experimental dataset has allowed us to make such an estimation. When defining the expected number of monolinks as the number of lysine, serine, threonine and tyrosine residues with a residue depth below 6.25Å (Figure S1), the data yields an average recovery rate of 50.64% (±14.06% standard deviation). Based on this we decided to implement a recovery rate of 50% when simulating monolinks.

To avoid using a residue depth cutoff to determine whether or not a residue can be monolinked in our calculations, we could potentially implement a probabilistic model for simulating monolinks (in a similar fashion to our probabilistic model we used to score crosslink length in MNXL (Matthew Allen Bullock et al., 2016)). This would entail weighting how likely a residue is to be monolinked based on its residue depth, given what is known from experimental data. The main impediment to this process is currently the lack of sufficient data to estimate such probabilities.

### Residue depth in MoDS

The exclusion of non-bulk water molecules from residue depth calculations was the main motivation to use this score, as a residue is much more likely to react with a crosslinking reagent if it is accessible to bulk water. Since solvent accessibility cannot measure this propensity accurately (Chakravarty and Varadarajan, 1999) we employed residue depth, which has been shown to be a better indicator of residue burial (due to its ability to distinguish between residues just below the protein surface and those closer to the core (Chakravarty and Varadarajan, 1999)). The complementary nature of residue depth and solvent accessibility (Liu et al., 2007; Tan et al., 2011, 2013), a property exploited in MNXL, may help explain the better performance of XLMO over MoDS/MNXL alone.

### Deep Residues in Experimental Models

When simulating monolinks, only residues with a residue depth below 6.25Å are selected as potential monolinks. This result generally fits our experimental data, except one example – X0987 – the poorest performing case in the experimental benchmark in terms of MoDS scoring. In this case, several of the monolinks which are found in the experimental dataset are very deep in the structure, for instance 45% of monolinks in this model have a residue depth greater than 6.25Å. The structure itself consists of two β-sandwich like domains connected by a short linker region, and most of these deep residues are present on the interface between the two domains. This could suggest that in solution, the two domains occupy a wider array of configurations with respect to one another, in comparison to the crystal, where they are likely to be more constrained. This is supported by the increased performance in terms of MoDS when the two domains are separated. It should also be noted that MNXL performance decreases when the domains are separated, possibly owing to the loss of several crosslinks between the two. This may suggest monolink information can be valuable in capturing dynamics of the protein and can give clues about which residues are located on the interface. Furthermore, this may also explain why X0987 was not modelled well in the Critical Assessment of Techniques for Protein Structure Prediction 13 (CASP13) (Kryshtafovych et al., 2019), where this model achieved a maximum GDT-TS score of 35.5% using an approach requiring no experimental information (Fajardo et al., 2019).

### Utility of MoDS

Some of the methods of covalent labeling used for XLMS use specific chemistry to target particular side chains. These methods typically target arginines, acidic side chains, cysteines, histidines, lysines, tryptophans and tyrosines (Mendoza and Vachet, 2009). Other methods that can be combined with MS utilize non-specific covalent modification. The most prevalent of these methods are oxidative radical footprinting, where photoactivated radicals label surface exposed residues (Cornwell et al., 2018), and Hydrodren Deuterium Exchange (HDX), where solvent accessible residues are modified with deuterium and can be identified by their rate of back exchange with hydrogen in water (Katta and Chait, 1993). MoDS should also be applicable to these methods, providing a score to evaluate a structural model using information from these covalent modifications.

### Experimental Considerations

One of the motivations for this study is that monolinks are produced as a by-product in an XLMS experiment. No deviation from existing protocols is required to produce them; however, the information they convey is not currently being utilised in the field. Furthermore, there are conditions where this monolink information could be more useful, such as when using shorter crosslinkers. Using shorter crosslinkers (e.g. BS2G, which has a 7.7Å spacer arm as opposed to the 11.4Å arm found in BS3/DSS) has been suggested to lead to fewer crosslinks being formed and a rise in the number of monolinks (Rozbeský et al., 2018). In these instances, including monolink information using MoDS is likely to be even more important for protein structure modelling as there will be fewer crosslink data points from which to draw useful information.

We show that monolinks can be used successfully in the modelling of protein structures. One type of information from an XLMS experiment, which has not been investigated is about the unmodified peptide, i.e an indication that a residue has not been modified by a crosslinker. However, since it is essential to perform a peptide enrichment step to remove many of these unmodified peptides, their disproportionate abundance with respect to crosslinks, monolinks and loop-links is unlikely to allowed their identification. Additionally, as described above, we cannot expect a 100% recovery rate for either crosslinks or monolinks. Therefore, we cannot assume that just because we have not detected a crosslink or monolink on a residue it cannot eliminate the theoretical possibility of that residue to chemically interact with the crosslinking reagent.

## SUMMARY

This study shows that, when used in conjunction with MNXL, our monolinks-based scoring function, MoDS, should improve the ranking of protein structure models. This is seen clearly in our simulated Benchmark. Our experimental dataset also shows, on average, a higher performance of the combined MNXL and MoDS scoring (XLMO), although this is inconclusive due to the the small size of the benchmark. To more accurately assess the effect of using experimentally obtained monolinks larger datasets would be required. Although there are many crosslink datasets in the literature, most of them do not provide information on peptides other than the crosslinks, making the investigation of monolinks difficult. Community wide deposition of whole datasets (e.g. PRIDE) would enhance our ability to make more interesting insights from this data.

## ACKNOWLEDGEMENTS

We thank Dr David Houldershaw for his technical help. We thank Dr Mark Williams and also members of the Topf and Thalassinos groups for helpful discussions, especially Dr Thomas Menneteau for his help with mass-spectrometry data processing. MT and KT are grateful for funding from the Wellcome Trust (209250/Z/17/Z).

## AUTHOR CONTRIBUTIONS

M.S. and M.T. designed the research. M.S carried out experiments. M.S, S.M., K.T., and M.S.M. analysed the data. M.S. and M.T. wrote the paper with contributions from all other authors.

## STAR METHODS

### Testing of MoDS on a Simulated Benchmark

For testing the scoring function, we used a large benchmark of protein structures and decoy models. This benchmark, which was downloaded from the Zhang Lab website, was generated using the 3DRobot decoy modelling algorithm (Deng et al., 2016). The benchmark contains 200 target PDB structures comprising a wide range of structural folds. Each of the 200 PDB structures have 300 decoy models associated with them, and they vary in similarity to the target structure (from 0 to ~12 Å Cα-RMSD).

Monolinks were simulated for each of the 200 target structures, based on known properties of the BS3 and DSS crosslinkers, *i.e.,* selecting only Lys, Thr, Ser, and Tyr. residue depth was calculated for each residue and those containing residue depth below 6.25Å were considered “true” simulated monolinks and therefore selected for further analysis. This 6.25Å cutoff was derived from an analysis performed on a smaller benchmark that we previously used (Matthew Allen Bullock et al., 2016), showing optimal performance (see STAR methods) when using 50-60% recovery rate (the percentage of experimentally observed crosslinked/monolink peptides, compared with the total theoretically possible ones), which is currently the estimate in the field (Figure S1). The simulated monolinks for each target structure were then applied to all corresponding decoy models in the dataset, and the models were then scored using MoDS (Figure S2).

### Testing the Effect of Monolink Recovery Rate

To investigate the effect of recovery on MoDS scoring we simulated monolinks at recovery rates for all integer percentage values from 1-100%. For each structure in the simulated benchmark we took the list of all potential monolinks (Lys, Thr, Ser, and Tyr residues in the target structure with a residue depth < 6.25Å) and selected a random set of amino acids for each given percentage recovery rate. This monolink list was then used to score all decoy structures with MoDS. Monolinks were randomly resampled 1000 times for each cutoff for each protein to gain a representative sample of different potential monolink configurations.

### Comparing MNXL and MoDS Scoring on Simulated Crosslinks and Monolinks

Model structures’ monolinks were simulated and randomly resampled with a constant recovery rate set at a 50%, which reflects the average recovery rate we see in our experimental dataset (see Discussion). Crosslinks were simulated at 20% recovery from a list of all possible Solvent Accessible Surface Distances (SASDs) using Jwalk (Bullock et al., 2018b; Matthew Allen Bullock et al., 2016). Since Jwalk calculates Cα-Cα SASDs, maximum distance of 33 Å was used to select “true” simulated crosslinks, to account for the length of the lysine side chains, the linker length and flexibility of the two (assuming the use of DSS and BS3 crosslinkers) (Bullock et al., 2018b; Matthew Allen Bullock et al., 2016).

Decoy models were scored with an *in-house* script that outputs both MNXL and MoDS (XLM tools available at: https://github.com/Topf-Lab/XLM-Tools). Scores are converted into Z-scores and a combination score (XLMO) is calculated by summing the two scores (Figure S2).

### Experimental Crosslink and Monolink Preparation

Experimental data containing both crosslink and monolink information was obtained from the Proteomic Identification Database (PRIDE) and was reprocessed (Martens et al., 2005) (Table 1). Reprocessing of the data was necessary for a fair comparison of the results and because often monolink data was not present in the associated publications. The criteria were: the protein was crosslinked by BS3/DSS, the protein was monomeric, separation of higher order oligomers was performed, either raw files or detailed CSV files including all peptides are provided and an experimentally derived PDB structure was available.

Raw files were converted to MGF files, which were then processed into mzXML files using msconvert (scan numbers were checked and corrected using an in-house script) (James et al., 2019). Fasta sequences were obtained for the PDB structures to be searched, and were used to generate reverse decoy sequences. These files were then used as input into the Xquest/Xprophet pipeline (Leitner et al., 2013), parameters for crosslink search were derived from the individual datasets. Xprophet crosslink assessment used an error range of −10 – 10 ppm (Leitner et al., 2013).

### Experimental Benchmark Decoy Model Generation

Benchmark decoy models were generated for assessing the performance of the scoring function. For each protein model 300 decoy structures were generated with a Cα-RMSD range of approximately 0-15Å. Two models were incomplete: CASP13 target Human Exonuclease V (X0975) (Cheng et al., 2019) and Creatine Kinase (Table 1) containing missing regions of 12 and 10 residues respectively. We used the *loopmodel* function in MODELLER9.22 (Webb and Sali, 2014) to complete these models. 100 models were generated and scored using SOAP potentials (Dong et al., 2013). The model with the best SOAP potential was selected (after manual inspection) as the “target” structure for the benchmark.

Next, random alignments for all benchmark cases were generated for the target sequence structure using an *in-house* script, which introduces random gaps in a constrained manner to ensure that they can produce model structures within an acceptable Cα-RMSD range. 300 sub-optimal alignments with gaps were produced and then used by MODELLER9.22 *Automodel* function (Webb and Sali, 2014) to output corresponding decoy models.

### Model Scoring and Performance Evaluation

A modified version of Jwalk (Matthew Allen Bullock et al., 2016) was produced, which allows Solvent Accessible Surface Distances (SASDs) to be calculated not only between Lys residues but also between Ser, Thr and Tyr residues. Jwalk was then applied to all decoy models per benchmark target. The DEPTH algorithm (Tan et al., 2011) was then run on these models to obtain residue depth information. Experimental recovery rates were calculated as the percentage of the experimental crosslinks and monolinks over the theoretical maximal number of crosslinks and monolinks, respectively.

Model structures were scored based on both crosslinks and monolinks, with crosslinks scored using MNXL (Matthew Allen Bullock et al., 2016) and monolinks using MoDS, and using the combined XLMO score. The relationship between MoDS and Cα-RMSD was evaluated using Spearman Correlation Coefficient, Precision, and the Area Under the ROC Curve (AUC) (Hand and Till, 2001). Precision is defined as TP/(TP+FP), where TP are True Positives and FP are False Positives, with a threshold of 4 Å and the top 10% scoring models (Table S1). ROC curves are calculated as the True Positive Rate (TPR) (Equation 1) against the True Negative Rate (TNR):

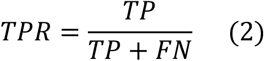

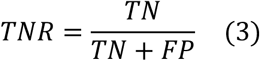

where FN are False Negatives and TN are True Negatives, and the threshold for the % top scoring models for deciding positive models being varied over the course of the calculation.

