## Supplemental Material for "Combining Information from Crosslinks and Monolinks in the Modelling of Protein Structures"

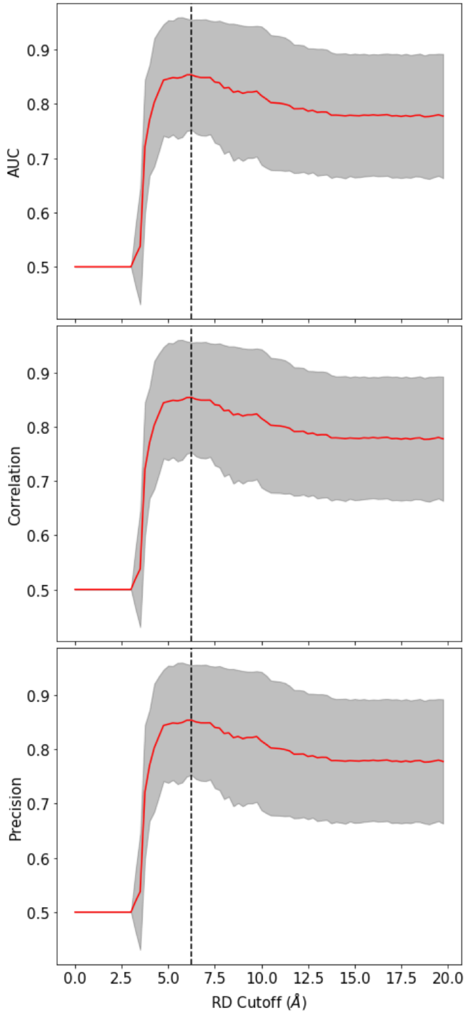


Figure S1: Plot showing MoDS performance when the maximum depth cutoff for simulating monolinks (i.e, identifying “true” monolinks) is varied. Monolinks were simulated at 50% recovery, red line shows mean performance over the benchmark from Bullock et al (Matthew Allen Bullock *et al.*, 2016), error bars show standard deviation. Black dotted line at 6.25Å shows the optimum performance.

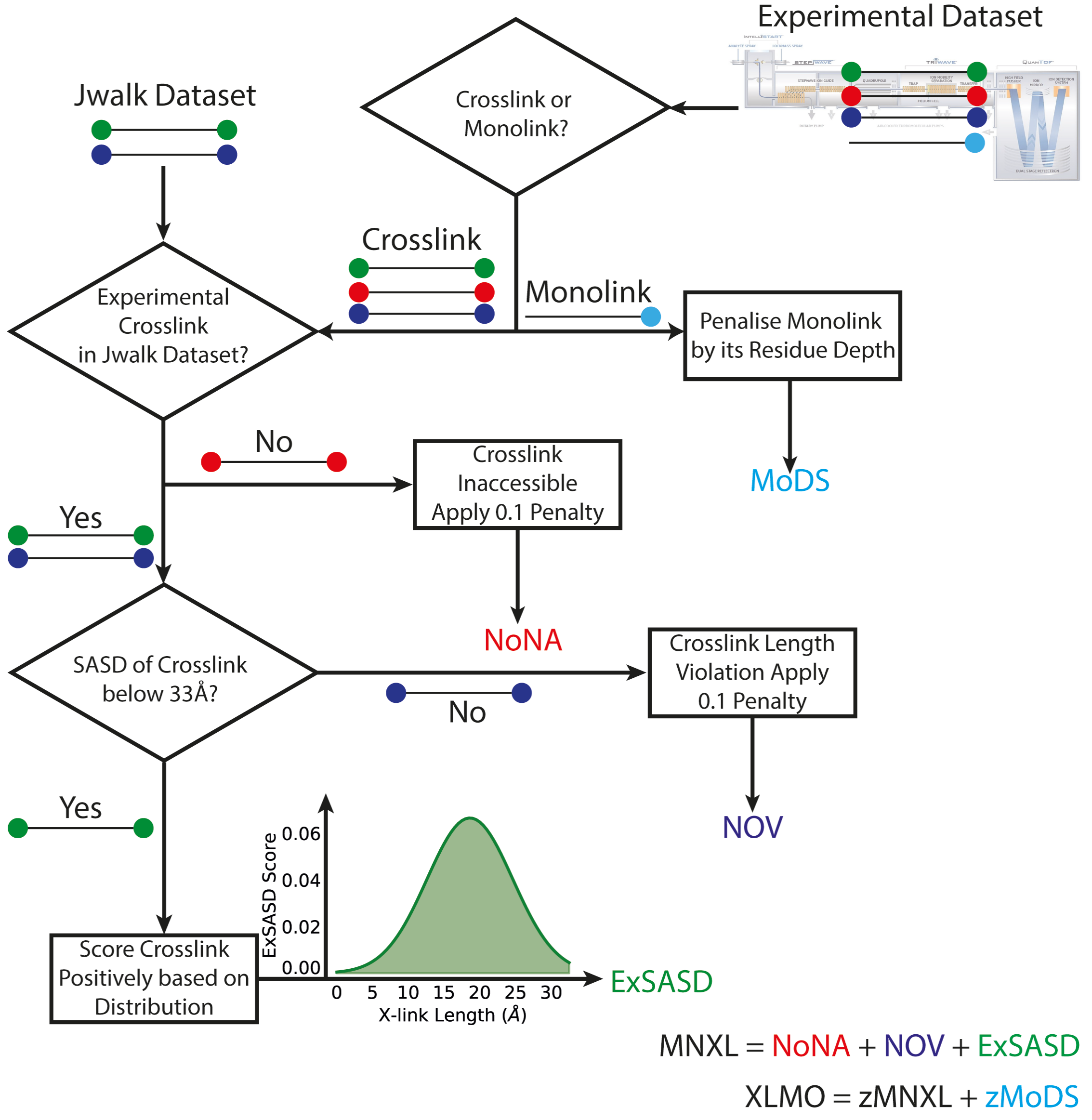


Figure S2: Flowchart showing the process for MNXL and XLMO scoring. Sticks represent crosslinkers, coloured circles show where a residue is attached

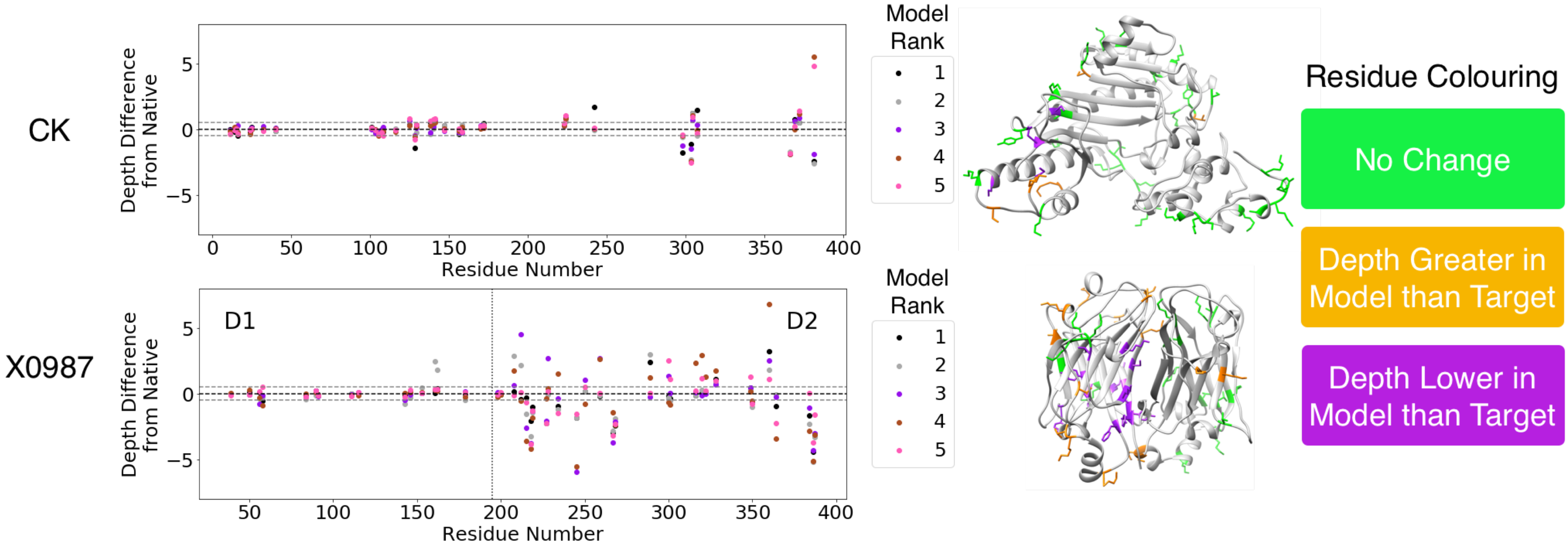


Figure S3: Plots showing the difference in RDs between monolinked residues in the native structure and in the 5 top-ranked models, based on MoDS (CK on top and X0987 in the bottom). Residues associated with difference > 0.5 Å or < -0.5 Å are shown on the structure, coloured red for depth decrease, green for depth increase and black for no change. For target structure X0987 d1 and d2 represent the residue range associated with domain 1 and domain 2, respectively.

**Table S1:** Definitions of binary classifiers.

| Value | Definition |
|---|---|
| TP | Cα-RMSD ≤ 4Å, above score threshold |
| FP | Cα-RMSD > 4Å, above score threshold |
| TN | Cα-RMSD > 4Å, below score threshold |
| FN | Cα-RMSD ≤ 4Å, below score threshold |

Table S2 - Statistics for the Simulated Benchmark

| | AUC | | | Correlation | | | Precision | | |
|---|---|---|---|---|---|---|---|---|---|
| Model | MNXL | MoDS | XLMO | MNXL | MoDS | XLMO | MNXL | MoDS | XLMO |
| 1WDDS | 0.50 | 0.88 | 0.88 | 0.00 | 0.66 | 0.66 | 0.27 | 0.66 | 0.66 |
| 3EINA | 0.90 | 0.87 | 0.92 | 0.81 | 0.75 | 0.84 | 0.82 | 0.73 | 0.85 |
| 3CHBD | 0.50 | 0.71 | 0.71 | 0.00 | 0.50 | 0.50 | 0.32 | 0.43 | 0.43 |
| 2X3MA | 0.98 | 0.91 | 0.99 | 0.57 | 0.42 | 0.54 | 0.94 | 0.69 | 0.93 |
| 2FHZB | 0.50 | 0.57 | 0.57 | 0.00 | 0.16 | 0.16 | 0.30 | 0.26 | 0.26 |
| 3R72A | 0.88 | 0.78 | 0.89 | 0.69 | 0.52 | 0.70 | 0.74 | 0.62 | 0.77 |
| 4IZXA | 0.76 | 0.73 | 0.79 | 0.57 | 0.50 | 0.62 | 0.53 | 0.39 | 0.54 |
| 4HKGA | 0.88 | 0.79 | 0.89 | 0.59 | 0.42 | 0.59 | 0.85 | 0.62 | 0.85 |
| 3WA4A | 0.75 | 0.71 | 0.78 | 0.44 | 0.39 | 0.50 | 0.53 | 0.43 | 0.57 |
| 3WDDA | 0.88 | 0.87 | 0.92 | 0.66 | 0.60 | 0.70 | 0.81 | 0.74 | 0.86 |
| 2J6BA | 0.88 | 0.81 | 0.89 | 0.68 | 0.52 | 0.68 | 0.85 | 0.70 | 0.87 |
| 4EGUA | 0.84 | 0.72 | 0.82 | 0.71 | 0.57 | 0.72 | 0.60 | 0.45 | 0.59 |
| 4GAIA | 0.84 | 0.70 | 0.84 | 0.61 | 0.50 | 0.68 | 0.80 | 0.39 | 0.75 |
| 2C2UA | 0.41 | 0.60 | 0.51 | 0.03 | 0.29 | 0.21 | 0.12 | 0.29 | 0.16 |
| 3WCQA | 0.91 | 0.80 | 0.91 | 0.72 | 0.51 | 0.71 | 0.85 | 0.60 | 0.84 |
| 1O7IA | 0.77 | 0.79 | 0.84 | 0.53 | 0.63 | 0.72 | 0.62 | 0.61 | 0.71 |
| 3MMHA | 0.78 | 0.67 | 0.77 | 0.60 | 0.29 | 0.54 | 0.56 | 0.43 | 0.58 |
| 2LISA | 0.91 | 0.86 | 0.93 | 0.74 | 0.57 | 0.74 | 0.82 | 0.70 | 0.86 |
| 3TEUA | 0.81 | 0.77 | 0.84 | 0.57 | 0.48 | 0.61 | 0.63 | 0.54 | 0.66 |
| 3WDNA | 0.84 | 0.87 | 0.90 | 0.66 | 0.67 | 0.75 | 0.72 | 0.75 | 0.82 |
| 3BY8A | 0.82 | 0.72 | 0.83 | 0.59 | 0.45 | 0.62 | 0.73 | 0.47 | 0.73 |
| 4G7XB | 0.50 | 0.83 | 0.83 | 0.00 | 0.51 | 0.51 | 0.35 | 0.66 | 0.66 |
| 4HQZA | 0.77 | 0.66 | 0.76 | 0.62 | 0.44 | 0.63 | 0.57 | 0.38 | 0.52 |
| 3IPJA | 0.85 | 0.67 | 0.81 | 0.72 | 0.42 | 0.66 | 0.70 | 0.34 | 0.60 |
| 4B97A | 0.79 | 0.80 | 0.85 | 0.61 | 0.61 | 0.71 | 0.64 | 0.67 | 0.72 |
| 1W0NA | 0.90 | 0.86 | 0.92 | 0.74 | 0.61 | 0.73 | 0.92 | 0.82 | 0.94 |
| 4OO4A | 0.89 | 0.72 | 0.85 | 0.77 | 0.49 | 0.73 | 0.75 | 0.45 | 0.67 |
| 2QJLA | 0.82 | 0.77 | 0.86 | 0.61 | 0.38 | 0.59 | 0.70 | 0.51 | 0.75 |
| 1BYIA | 0.87 | 0.85 | 0.90 | 0.73 | 0.70 | 0.78 | 0.76 | 0.67 | 0.78 |
| 3QZXA | 0.76 | 0.78 | 0.82 | 0.56 | 0.51 | 0.64 | 0.54 | 0.49 | 0.60 |
| 3ZSJA | 0.81 | 0.82 | 0.86 | 0.61 | 0.60 | 0.68 | 0.73 | 0.66 | 0.78 |
| 1F86A | 0.85 | 0.81 | 0.88 | 0.67 | 0.56 | 0.71 | 0.69 | 0.58 | 0.73 |
| 3ULJA | 0.71 | 0.75 | 0.78 | 0.40 | 0.39 | 0.47 | 0.54 | 0.45 | 0.56 |
| 1E5KA | 0.77 | 0.76 | 0.81 | 0.51 | 0.46 | 0.58 | 0.59 | 0.53 | 0.64 |
| 4HRRB | 0.50 | 0.84 | 0.84 | 0.00 | 0.64 | 0.64 | 0.34 | 0.74 | 0.74 |
| 3RHBA | 0.87 | 0.73 | 0.86 | 0.73 | 0.53 | 0.74 | 0.82 | 0.51 | 0.79 |
| 4CO8A | 0.60 | 0.54 | 0.60 | 0.27 | 0.11 | 0.25 | 0.39 | 0.18 | 0.33 |
| 4MZIA | 0.86 | 0.83 | 0.89 | 0.60 | 0.65 | 0.72 | 0.73 | 0.61 | 0.76 |
| 4I4OA | 0.83 | 0.91 | 0.89 | 0.73 | 0.76 | 0.80 | 0.59 | 0.74 | 0.72 |
| 1R29A | 0.75 | 0.74 | 0.79 | 0.40 | 0.35 | 0.44 | 0.54 | 0.44 | 0.54 |
| 3NJNA | 0.77 | 0.61 | 0.74 | 0.43 | 0.19 | 0.38 | 0.60 | 0.26 | 0.53 |
| 3ZJAA | 0.97 | 0.90 | 0.97 | 0.70 | 0.55 | 0.67 | 0.95 | 0.77 | 0.95 |
| 4HE6A | 0.75 | 0.76 | 0.80 | 0.39 | 0.40 | 0.47 | 0.62 | 0.55 | 0.66 |

| | | | | | | | | | |
|---|---|---|---|---|---|---|---|---|---|
| 4A7UA | 0.82 | 0.88 | 0.91 | 0.56 | 0.64 | 0.71 | 0.62 | 0.73 | 0.79 |
| 1TU9A | 0.72 | 0.67 | 0.74 | 0.37 | 0.29 | 0.40 | 0.59 | 0.39 | 0.57 |
| 4NBPA | 0.64 | 0.55 | 0.62 | 0.42 | 0.29 | 0.42 | 0.36 | 0.18 | 0.29 |
| 1R8SA | 0.81 | 0.77 | 0.84 | 0.53 | 0.53 | 0.61 | 0.69 | 0.60 | 0.72 |
| 4EICA | 0.78 | 0.52 | 0.69 | 0.46 | 0.05 | 0.32 | 0.63 | 0.19 | 0.45 |
| 2OVJA | 0.86 | 0.77 | 0.86 | 0.72 | 0.53 | 0.72 | 0.74 | 0.59 | 0.76 |
| 4DQJA | 0.74 | 0.75 | 0.80 | 0.47 | 0.49 | 0.59 | 0.63 | 0.51 | 0.67 |
| 1NG6A | 0.96 | 0.87 | 0.97 | 0.69 | 0.50 | 0.68 | 0.83 | 0.56 | 0.86 |
| 1PSRA | 0.81 | 0.83 | 0.87 | 0.54 | 0.41 | 0.55 | 0.58 | 0.60 | 0.70 |
| 3NE0A | 0.81 | 0.81 | 0.85 | 0.67 | 0.68 | 0.77 | 0.69 | 0.64 | 0.74 |
| 1WN2A | 0.85 | 0.80 | 0.87 | 0.69 | 0.59 | 0.72 | 0.71 | 0.57 | 0.73 |
| 3D32A | 0.78 | 0.73 | 0.80 | 0.55 | 0.46 | 0.58 | 0.56 | 0.41 | 0.55 |
| 2OHWA | 0.91 | 0.87 | 0.94 | 0.64 | 0.54 | 0.67 | 0.81 | 0.70 | 0.86 |
| 2HS1A | 0.71 | 0.69 | 0.74 | 0.49 | 0.52 | 0.60 | 0.49 | 0.39 | 0.50 |
| 2PIEA | 0.65 | 0.76 | 0.74 | 0.37 | 0.49 | 0.52 | 0.37 | 0.48 | 0.46 |
| 3FYMA | 0.70 | 0.52 | 0.65 | 0.42 | 0.09 | 0.34 | 0.51 | 0.16 | 0.38 |
| 1FK5A | 0.77 | 0.64 | 0.74 | 0.60 | 0.20 | 0.48 | 0.62 | 0.45 | 0.64 |
| 4JEAA | 0.68 | 0.51 | 0.63 | 0.39 | 0.03 | 0.28 | 0.56 | 0.23 | 0.45 |
| 3IX3A | 0.82 | 0.83 | 0.87 | 0.56 | 0.55 | 0.64 | 0.73 | 0.69 | 0.78 |
| 3N01A | 0.80 | 0.62 | 0.78 | 0.56 | 0.25 | 0.54 | 0.73 | 0.32 | 0.60 |
| 4HVYA | 0.83 | 0.82 | 0.87 | 0.61 | 0.61 | 0.69 | 0.70 | 0.60 | 0.75 |
| 3OGNA | 0.77 | 0.88 | 0.88 | 0.55 | 0.67 | 0.70 | 0.55 | 0.74 | 0.72 |
| 2XQQA | 0.89 | 0.75 | 0.87 | 0.63 | 0.33 | 0.55 | 0.79 | 0.43 | 0.72 |
| 1CY5A | 0.71 | 0.64 | 0.71 | 0.41 | 0.31 | 0.42 | 0.49 | 0.37 | 0.47 |
| 2WF7A | 0.59 | 0.82 | 0.75 | 0.19 | 0.58 | 0.47 | 0.40 | 0.68 | 0.58 |
| 4J1PA | 0.80 | 0.72 | 0.81 | 0.56 | 0.38 | 0.55 | 0.72 | 0.48 | 0.68 |
| 4BMBA | 0.71 | 0.68 | 0.74 | 0.50 | 0.45 | 0.56 | 0.58 | 0.46 | 0.57 |
| 2Y6HA | 0.80 | 0.84 | 0.87 | 0.54 | 0.65 | 0.68 | 0.65 | 0.64 | 0.71 |
| 3W42A | 0.80 | 0.79 | 0.83 | 0.60 | 0.56 | 0.64 | 0.62 | 0.62 | 0.69 |
| 3A38A | 0.62 | 0.71 | 0.70 | 0.30 | 0.37 | 0.42 | 0.35 | 0.39 | 0.40 |
| 1T6UA | 0.69 | 0.56 | 0.66 | 0.34 | 0.12 | 0.29 | 0.49 | 0.25 | 0.44 |
| 3K67A | 0.88 | 0.82 | 0.89 | 0.67 | 0.58 | 0.69 | 0.79 | 0.68 | 0.82 |
| 3T3LA | 0.82 | 0.78 | 0.84 | 0.67 | 0.55 | 0.69 | 0.64 | 0.53 | 0.64 |
| 2NSZA | 0.93 | 0.84 | 0.94 | 0.76 | 0.58 | 0.76 | 0.92 | 0.64 | 0.90 |
| 4I2UA | 0.88 | 0.77 | 0.88 | 0.75 | 0.57 | 0.75 | 0.78 | 0.51 | 0.76 |
| 2FQ3A | 0.89 | 0.81 | 0.91 | 0.64 | 0.48 | 0.65 | 0.78 | 0.58 | 0.81 |
| 3WDCA | 0.83 | 0.88 | 0.90 | 0.55 | 0.66 | 0.69 | 0.76 | 0.78 | 0.86 |
| 3CI3A | 0.73 | 0.73 | 0.76 | 0.55 | 0.58 | 0.63 | 0.45 | 0.40 | 0.46 |
| 4MAXA | 0.74 | 0.81 | 0.82 | 0.53 | 0.56 | 0.63 | 0.52 | 0.64 | 0.63 |
| 4H7WA | 0.94 | 0.92 | 0.95 | 0.82 | 0.82 | 0.87 | 0.90 | 0.84 | 0.94 |
| 3PIWA | 0.86 | 0.80 | 0.87 | 0.64 | 0.56 | 0.67 | 0.71 | 0.61 | 0.73 |
| 2IMFA | 0.76 | 0.84 | 0.85 | 0.53 | 0.61 | 0.67 | 0.65 | 0.75 | 0.77 |
| 4B62A | 0.81 | 0.72 | 0.82 | 0.60 | 0.37 | 0.59 | 0.69 | 0.47 | 0.68 |
| 4GZCA | 0.89 | 0.84 | 0.91 | 0.72 | 0.62 | 0.74 | 0.74 | 0.65 | 0.77 |
| 1NWZA | 0.83 | 0.83 | 0.87 | 0.58 | 0.59 | 0.67 | 0.67 | 0.64 | 0.74 |
| 3RZNA | 0.85 | 0.83 | 0.87 | 0.74 | 0.66 | 0.75 | 0.70 | 0.62 | 0.70 |

| | | | | | | | | | |
|---|---|---|---|---|---|---|---|---|---|
| 1Q5YA | 0.70 | 0.58 | 0.69 | 0.41 | 0.13 | 0.36 | 0.57 | 0.28 | 0.52 |
| 1GP0A | 0.78 | 0.84 | 0.85 | 0.46 | 0.54 | 0.57 | 0.69 | 0.71 | 0.79 |
| 2FHZA | 0.88 | 0.84 | 0.89 | 0.75 | 0.65 | 0.76 | 0.71 | 0.62 | 0.70 |
| 1NKIA | 0.89 | 0.89 | 0.93 | 0.57 | 0.61 | 0.65 | 0.74 | 0.71 | 0.79 |
| 3L9AX | 0.50 | 0.76 | 0.76 | 0.00 | 0.41 | 0.41 | 0.29 | 0.58 | 0.58 |
| 4B9CA | 0.81 | 0.86 | 0.88 | 0.60 | 0.70 | 0.74 | 0.60 | 0.70 | 0.72 |
| 3H87A | 0.93 | 0.86 | 0.94 | 0.78 | 0.69 | 0.82 | 0.89 | 0.70 | 0.90 |
| 3L42A | 0.81 | 0.78 | 0.84 | 0.50 | 0.43 | 0.54 | 0.63 | 0.47 | 0.63 |
| 1C5EA | 0.92 | 0.87 | 0.94 | 0.80 | 0.68 | 0.82 | 0.86 | 0.70 | 0.88 |
| 256BA | 0.84 | 0.63 | 0.79 | 0.64 | 0.26 | 0.55 | 0.77 | 0.40 | 0.72 |
| 4DRIB | 0.50 | 0.51 | 0.51 | 0.00 | 0.03 | 0.03 | 0.34 | 0.20 | 0.20 |
| 4BK7A | 0.83 | 0.78 | 0.86 | 0.51 | 0.48 | 0.58 | 0.70 | 0.50 | 0.71 |
| 3D3BA | 0.84 | 0.77 | 0.85 | 0.57 | 0.53 | 0.64 | 0.72 | 0.45 | 0.68 |
| 4ENFA | 0.95 | 0.94 | 0.97 | 0.64 | 0.64 | 0.67 | 0.90 | 0.87 | 0.92 |
| 2J9WA | 0.66 | 0.64 | 0.68 | 0.37 | 0.29 | 0.40 | 0.48 | 0.42 | 0.53 |
| 3FKEA | 0.84 | 0.64 | 0.80 | 0.65 | 0.26 | 0.56 | 0.72 | 0.27 | 0.60 |
| 1MF7A | 0.89 | 0.78 | 0.90 | 0.65 | 0.40 | 0.63 | 0.83 | 0.56 | 0.82 |
| 2FCJA | 0.85 | 0.70 | 0.85 | 0.60 | 0.35 | 0.59 | 0.72 | 0.38 | 0.70 |
| 3ZOOA | 0.91 | 0.82 | 0.90 | 0.69 | 0.52 | 0.67 | 0.88 | 0.66 | 0.88 |
| 4FP5D | 0.50 | 0.74 | 0.74 | 0.00 | 0.44 | 0.44 | 0.34 | 0.46 | 0.46 |
| 3AIAA | 0.93 | 0.84 | 0.94 | 0.72 | 0.55 | 0.72 | 0.94 | 0.72 | 0.93 |
| 2BMOB | 0.50 | 0.75 | 0.75 | 0.00 | 0.53 | 0.53 | 0.28 | 0.50 | 0.50 |
| 1WPUA | 0.85 | 0.80 | 0.88 | 0.70 | 0.61 | 0.74 | 0.77 | 0.57 | 0.77 |
| 2QFEA | 0.87 | 0.61 | 0.80 | 0.62 | 0.28 | 0.55 | 0.69 | 0.20 | 0.54 |
| 2WDSA | 0.46 | 0.79 | 0.74 | 0.03 | 0.56 | 0.49 | 0.31 | 0.56 | 0.53 |
| 3PLWA | 0.75 | 0.62 | 0.75 | 0.44 | 0.23 | 0.44 | 0.59 | 0.23 | 0.54 |
| 4A56A | 0.67 | 0.67 | 0.72 | 0.29 | 0.26 | 0.35 | 0.50 | 0.36 | 0.51 |
| 2XU3A | 0.82 | 0.88 | 0.89 | 0.68 | 0.68 | 0.76 | 0.64 | 0.76 | 0.77 |
| 3VZ9B | 0.50 | 0.71 | 0.71 | 0.00 | 0.45 | 0.45 | 0.34 | 0.39 | 0.39 |
| 4C45A | 0.83 | 0.78 | 0.86 | 0.45 | 0.32 | 0.45 | 0.73 | 0.60 | 0.77 |
| 2FVVA | 0.76 | 0.69 | 0.77 | 0.52 | 0.52 | 0.61 | 0.57 | 0.36 | 0.56 |
| 4KZVA | 0.78 | 0.72 | 0.78 | 0.65 | 0.56 | 0.67 | 0.60 | 0.46 | 0.58 |
| 1N8VA | 0.78 | 0.62 | 0.74 | 0.57 | 0.23 | 0.48 | 0.67 | 0.41 | 0.61 |
| 1V4PA | 0.91 | 0.94 | 0.95 | 0.75 | 0.72 | 0.78 | 0.89 | 0.87 | 0.91 |
| 4E3YA | 0.79 | 0.87 | 0.84 | 0.70 | 0.79 | 0.78 | 0.60 | 0.74 | 0.69 |
| 1ZI8A | 0.82 | 0.80 | 0.85 | 0.68 | 0.60 | 0.71 | 0.69 | 0.58 | 0.70 |
| 3SY1A | 0.88 | 0.89 | 0.93 | 0.56 | 0.55 | 0.62 | 0.83 | 0.77 | 0.89 |
| 4NJYA | 0.70 | 0.71 | 0.75 | 0.41 | 0.37 | 0.48 | 0.47 | 0.47 | 0.54 |
| 2QCPX | 0.50 | 0.75 | 0.75 | 0.00 | 0.37 | 0.37 | 0.28 | 0.41 | 0.41 |
| 3A6RA | 0.71 | 0.68 | 0.73 | 0.43 | 0.34 | 0.47 | 0.55 | 0.43 | 0.57 |
| 2BT9A | 0.90 | 0.80 | 0.90 | 0.71 | 0.49 | 0.69 | 0.84 | 0.61 | 0.84 |
| 4IL7A | 0.72 | 0.72 | 0.75 | 0.42 | 0.29 | 0.41 | 0.55 | 0.48 | 0.55 |
| 3NOQA | 0.88 | 0.82 | 0.91 | 0.49 | 0.43 | 0.54 | 0.80 | 0.59 | 0.82 |
| 2WNPF | 0.50 | 0.85 | 0.85 | 0.00 | 0.76 | 0.76 | 0.38 | 0.71 | 0.71 |
| 4GMQA | 0.76 | 0.70 | 0.78 | 0.56 | 0.41 | 0.58 | 0.55 | 0.42 | 0.55 |
| 1TJXA | 0.74 | 0.83 | 0.84 | 0.48 | 0.59 | 0.64 | 0.56 | 0.68 | 0.72 |

| | | | | | | | | | |
|---|---|---|---|---|---|---|---|---|---|
| 2XODA | 0.78 | 0.68 | 0.77 | 0.59 | 0.41 | 0.58 | 0.72 | 0.45 | 0.68 |
| 4J74A | 0.86 | 0.81 | 0.88 | 0.70 | 0.59 | 0.73 | 0.78 | 0.57 | 0.77 |
| 4F1WA | 0.82 | 0.86 | 0.88 | 0.60 | 0.69 | 0.73 | 0.77 | 0.75 | 0.85 |
| 1YPQA | 0.78 | 0.79 | 0.83 | 0.59 | 0.64 | 0.71 | 0.59 | 0.59 | 0.68 |
| 1MUNA | 0.88 | 0.74 | 0.87 | 0.73 | 0.55 | 0.74 | 0.75 | 0.47 | 0.71 |
| 1G8AA | 0.82 | 0.84 | 0.86 | 0.65 | 0.66 | 0.72 | 0.64 | 0.63 | 0.68 |
| 3IE4A | 0.73 | 0.67 | 0.75 | 0.45 | 0.31 | 0.47 | 0.49 | 0.38 | 0.53 |
| 3HNYM | 0.50 | 0.76 | 0.76 | 0.00 | 0.56 | 0.56 | 0.30 | 0.47 | 0.47 |
| 2O9UX | 0.50 | 0.47 | 0.47 | 0.00 | -0.13 | -0.13 | 0.30 | 0.12 | 0.12 |
| 3PMCA | 0.86 | 0.70 | 0.83 | 0.72 | 0.43 | 0.66 | 0.79 | 0.48 | 0.73 |
| 3CCDA | 0.67 | 0.53 | 0.61 | 0.34 | 0.03 | 0.18 | 0.40 | 0.15 | 0.25 |
| 3IVVA | 0.89 | 0.86 | 0.91 | 0.75 | 0.67 | 0.77 | 0.81 | 0.69 | 0.83 |
| 4FTFA | 0.82 | 0.69 | 0.82 | 0.50 | 0.14 | 0.40 | 0.72 | 0.34 | 0.63 |
| 2JLIA | 0.77 | 0.73 | 0.80 | 0.56 | 0.44 | 0.58 | 0.66 | 0.48 | 0.65 |
| 3ZRXA | 0.74 | 0.50 | 0.64 | 0.25 | -0.17 | 0.05 | 0.61 | 0.13 | 0.35 |
| 4N5UA | 0.64 | 0.65 | 0.68 | 0.20 | 0.16 | 0.23 | 0.45 | 0.37 | 0.44 |
| 3CIMA | 0.81 | 0.80 | 0.85 | 0.55 | 0.54 | 0.62 | 0.75 | 0.68 | 0.82 |
| 2C4JA | 0.89 | 0.88 | 0.92 | 0.81 | 0.77 | 0.85 | 0.78 | 0.76 | 0.83 |
| 2O90A | 0.47 | 0.73 | 0.61 | -0.03 | 0.40 | 0.22 | 0.28 | 0.51 | 0.40 |
| 3VUBA | 0.83 | 0.66 | 0.81 | 0.59 | 0.28 | 0.55 | 0.69 | 0.32 | 0.60 |
| 3NBCA | 0.88 | 0.86 | 0.91 | 0.80 | 0.69 | 0.81 | 0.73 | 0.68 | 0.78 |
| 1FX2A | 0.77 | 0.68 | 0.75 | 0.67 | 0.60 | 0.68 | 0.49 | 0.28 | 0.41 |
| 4JF8A | 0.79 | 0.83 | 0.85 | 0.62 | 0.68 | 0.73 | 0.61 | 0.61 | 0.70 |
| 4LR6A | 0.88 | 0.88 | 0.92 | 0.61 | 0.58 | 0.67 | 0.76 | 0.68 | 0.82 |
| 4EETB | 0.50 | 0.71 | 0.71 | 0.00 | 0.27 | 0.27 | 0.29 | 0.38 | 0.38 |
| 1UKFA | 0.97 | 0.95 | 0.99 | 0.70 | 0.63 | 0.71 | 0.89 | 0.82 | 0.94 |
| 1YN3A | 0.93 | 0.89 | 0.95 | 0.76 | 0.67 | 0.79 | 0.89 | 0.77 | 0.93 |
| 2YOIA | 0.85 | 0.75 | 0.86 | 0.69 | 0.44 | 0.67 | 0.80 | 0.54 | 0.78 |
| 3WDEA | 0.84 | 0.90 | 0.91 | 0.61 | 0.68 | 0.71 | 0.75 | 0.80 | 0.85 |
| 2ZNRA | 0.92 | 0.89 | 0.94 | 0.76 | 0.63 | 0.76 | 0.84 | 0.76 | 0.88 |
| 1R6JA | 0.62 | 0.77 | 0.79 | 0.22 | 0.46 | 0.49 | 0.37 | 0.51 | 0.55 |
| 3PUCA | 0.92 | 0.78 | 0.90 | 0.69 | 0.46 | 0.65 | 0.86 | 0.54 | 0.79 |
| 4AQOA | 0.81 | 0.71 | 0.82 | 0.61 | 0.37 | 0.59 | 0.66 | 0.48 | 0.66 |
| 1N13B | 0.50 | 0.88 | 0.88 | 0.00 | 0.71 | 0.71 | 0.32 | 0.68 | 0.68 |
| 1XT5A | 0.84 | 0.69 | 0.82 | 0.42 | 0.23 | 0.38 | 0.65 | 0.37 | 0.58 |
| 4M2PA | 0.84 | 0.83 | 0.87 | 0.74 | 0.69 | 0.78 | 0.69 | 0.61 | 0.72 |
| 1Q92A | 0.90 | 0.85 | 0.91 | 0.53 | 0.49 | 0.56 | 0.81 | 0.67 | 0.82 |
| 2HEWF | 0.50 | 0.87 | 0.87 | 0.00 | 0.71 | 0.71 | 0.34 | 0.71 | 0.71 |
| 1SAUA | 0.83 | 0.81 | 0.85 | 0.68 | 0.65 | 0.73 | 0.65 | 0.58 | 0.67 |
| 1MY7A | 0.73 | 0.71 | 0.77 | 0.49 | 0.35 | 0.52 | 0.55 | 0.41 | 0.58 |
| 2ACFA | 0.84 | 0.82 | 0.87 | 0.70 | 0.58 | 0.72 | 0.70 | 0.68 | 0.77 |
| 2COVD | 0.50 | 0.73 | 0.73 | 0.00 | 0.46 | 0.46 | 0.34 | 0.45 | 0.45 |
| 2VZCA | 0.81 | 0.81 | 0.85 | 0.57 | 0.57 | 0.65 | 0.66 | 0.60 | 0.69 |
| 2YH5A | 0.95 | 0.90 | 0.97 | 0.76 | 0.59 | 0.76 | 0.80 | 0.74 | 0.89 |
| 4JFIA | 0.82 | 0.66 | 0.79 | 0.57 | 0.27 | 0.50 | 0.64 | 0.29 | 0.54 |
| 2CCVA | 0.86 | 0.68 | 0.83 | 0.69 | 0.36 | 0.63 | 0.80 | 0.43 | 0.70 |

| | | | | | | | | | |
|---|---|---|---|---|---|---|---|---|---|
| 3O7BA | 0.82 | 0.90 | 0.91 | 0.59 | 0.49 | 0.62 | 0.46 | 0.59 | 0.60 |
| 3TOWA | 0.97 | 0.96 | 0.99 | 0.71 | 0.67 | 0.73 | 0.95 | 0.91 | 0.98 |
| 3M9QA | 0.86 | 0.69 | 0.86 | 0.54 | 0.19 | 0.47 | 0.71 | 0.38 | 0.69 |
| 3SO6A | 0.80 | 0.85 | 0.88 | 0.61 | 0.59 | 0.68 | 0.65 | 0.70 | 0.76 |
| 2J73A | 0.81 | 0.72 | 0.82 | 0.57 | 0.36 | 0.55 | 0.72 | 0.46 | 0.71 |
| Mean | 0.78 | 0.77 | 0.82 | 0.52 | 0.48 | 0.59 | 0.64 | 0.54 | 0.67 |
| STD | 0.13 | 0.10 | 0.09 | 0.22 | 0.18 | 0.16 | 0.17 | 0.17 | 0.17 |
